# Taking aim at the perceptual side of motor learning: Exploring how explicit and implicit learning encode perceptual error information through depth vision

**DOI:** 10.1101/2021.04.02.438242

**Authors:** Carlo Campagnoli, Fulvio Domini, Jordan A. Taylor

## Abstract

Motor learning in visuomotor adaptation tasks results from both explicit and implicit processes, each responding differently to an error signal. While the motor output side of these processes is extensively studied, their visual input side is relatively unknown. We investigated if and how depth perception affects the computation of error information by explicit and implicit motor learning. Two groups of participants threw virtual darts at a virtual dartboard while receiving perturbed endpoint feedback. The Delayed group was allowed to re-aim and their feedback was delayed to emphasize explicit learning, while the Clamped group received clamped cursor feedback which they were told to ignore, and continued to aim straight at the target to emphasize implicit adaptation. Both groups played this game in a highly detailed virtual environment (Depth condition) and in an empty environment (No-Depth condition). The Delayed group showed an increase in error sensitivity under Depth relative to No-Depth conditions. In contrast, the Clamped group adapted to the same degree under both conditions. The movement kinematics of the Delayed participants also changed under the Depth condition, consistent with the target appearing more distant, unlike the Clamped group. A comparison of the Delayed behavioral data with a perceptual task from the same individuals showed that the effect of the Depth condition on the re-aiming direction was consistent with an increase in the scaling of the error distance and size. These findings suggest that explicit and implicit learning processes may rely on different sources of perceptual information.

**New & Noteworthy:** We leveraged a classic sensorimotor adaptation task to perform a first systematic assessment of the role of perceptual cues in the estimation of an error signal in the 3D space during motor learning. We crossed two conditions presenting different amounts of depth information, with two manipulations emphasizing explicit and implicit learning processes. Explicit learning responded to the visual conditions, consistent with perceptual reports, while implicit learning appeared to be independent of them.

## Introduction

Adaptation is a fundamental process that helps maintain calibration of the motor system in a constantly changing environment. Understanding this process has been a central focus of numerous studies over the past twenty years, seeking to characterize the error sensitivity of the motor system. Typically, these studies have adopted a system-identification approach to model how humans adjust their motor behavior in response to experimental perturbations of visual feedback, such as in prism adaptation (Hanajima et al., 2015; Petitet et al., 2018) and visuomotor rotations (Wei and Kording, 2010; Semrau et al., 2012; Marko et al., 2012; Hutter and Taylor, 2018). Indeed, this work has led to elegant computational models that can account for the trial-by-trial output of the motor system in response to errors (Thoroughman and Shadmehr, 2000; Kording and Wolpert, 2004; Cheng and Sabes, 2006; He et al., 2016).

Conversely, the input side of the adaptation process remains less explored. One aspect of this issue concerns how visual error information gets encoded for motor learning in the first place. It is often tacitly assumed that the visual input is transformed into an accurate representation of the physical scene, which seems at odds with the fact that visual perception is frequently unreliable and biased (Goldstein and Brockmole, 2016). There are practical reasons that may have favored the assumption that visual representations are metric and veridical in the context of this research. Most visuomotor adaptation studies have been constrained to a 2D plane, such as that of a monitor or a flat frontoparallel surface, to display feedback of reaching (i.e., rotations). Although pioneering studies on prism adaptation involved actions that were actually embedded in the three-dimensional space, such as pointing or throwing, they typically required knowledge of visual direction alone, while the distance to the targets was kept fixed without systematic manipulation or control of the depth cues in the environment (Redding and Wallace, 1988, 1993).

An advantage of operating in a simplified space is that any potential discrepancies between the visual representation and the physical world are likely to be quite small. However, most everyday life actions take place in a wider and three-dimensional environment, where the relationship between the visual input and the physical world cannot be determined as straightforwardly. To solve the problem of best interpreting the 3D layout of the scene (or to attempt to), a variety of visual cues must be integrated together to generate an estimate. Even when a simple line is seen on a rotated monitor or from an angle, it projects differently onto each eye, such that the estimation of its length or orientation requires the combination of multiple visual signals, both retinal and extraretinal (Howard and Rogers, 1995). In fact, the integration of visual cues is necessary for the estimation of almost any depth property. For example, the distance between two points on a screen, one representing a target and the other a cursor, cannot be quantified metrically, presumably to plan an accurate motor correction, unless it is scaled by an estimate of the viewing distance (Howard, 2002). Integrating depth information accurately is therefore critically important when reaching in 3D space, both in terms of motor planning and error correction. This unveils an interesting puzzle.

First off, depth vision is far from being accurate and, in fact, it often leads to perceptual distortions of space. Depth estimates fluctuate with several factors, including the type of task (Glennerster et al., 1996; Bradshaw et al., 2000), the viewing distance (Johnston, 1991; Campagnoli et al., 2017), the type of depth cues available (Campagnoli and Domini, 2019) and even following sensorimotor adaptation itself (Volcic et al., 2013). These observations raise the question as to what information the motor system relies on when scaling error size information to adjust to visual perturbations in a 3D environment.

Secondly, motor learning is not a unitary process. Sensorimotor adaptation of goal-directed actions is known to be shaped by the interplay between explicit re-aiming and implicit motor adaptation processes (Taylor and Ivry, 2012; Huberdeau et al., 2015; McDougle et al., 2016). While both can be defined as error-minimization processes, numerous findings indicate that each process operates on different types of error information. Explicit re-aiming is thought to be responsive to task errors (i.e., whether the target was hit or missed), resulting in a change in the intended aiming location (Taylor and Ivry, 2011; Leow et al., 2016; Morehead et al., 2017; Butcher and Taylor, 2018). Importantly, it is under volitional control, highly flexible to task demands, and sensitive to contextual information (Bond and Taylor, 2015, 2017; Poh and Taylor, 2019; Schween et al., 2018, 2019). Implicit adaptation, on the other hand, is triggered by sensory-prediction errors (a mismatch between the expected sensory outcome of a movement and the actual sensory feedback), resulting in an update to a sensorimotor mapping (Mazzoni and Krakauer, 2006; Morehead et al., 2017; Kim et al., 2018). Unlike explicit aiming, implicit adaptation appears to be obligatory, displays a stereotyped output, and evidence for contextual sensitivity is scant (Mazzoni and Krakauer, 2006; Bond and Taylor, 2015; Morehead et al., 2017; Kim et al., 2018; Schween et al., 2018, 2019). Both processes also appear to be reliant on different neural circuits. Explicit re-aiming is likely supported by neural networks associated with working memory, decision making, and planning, such as the prefrontal and parietal cortices (Slachevsky et al., 2001, 2003; Taylor and Ivry, 2014; McDougle and Taylor, 2019; de Brouwer et al., 2018; Anguera et al 2010; Christou et al 2016; Holland et al., 2019). Implicit adaptation has been repeatedly linked with the function of the cerebellum and may represent the updating of an internal forward model (Miall and Wolpert 1996; Martin and Thach, 1996; Tseng et al., 2007; Taylor and Ivry, 2011; Schlerf et al., 2013; Butcher et al., 2017).

Here we set out to explore if and how explicit and implicit processes incorporate multiple sources of depth information during sensorimotor adaptation, through a 3D version of a visuomotor rotation task. To have effective control of nearly all the depth cues in the scene, participants performed the task in virtual reality. They were asked to simulate “throwing” a virtual dart at a virtual dartboard positioned at varying distances in front of them by making horizontal, planar reaching movements on a tabletop. To determine the potential influence of visual depth cues on explicit and implicit learning, the experiment had a 2×2 factorial design contrasting two viewing conditions against two types of endpoint feedback. For the viewing conditions, we tested a Depth condition, where participants played darts in a full scale “tavern” scene, as rich in depth cues as there would be in the real world (perspective, texture, stereo disparity, objects of familiar size, et cetera), and a No-Depth condition, where participants played darts in a nondescript gray environment in which the dartboard was the only object visible. As for the type of feedback, one group of participants performed the task while receiving delayed endpoint feedback, which appears to weaken implicit adaptation and require explicit learning, whereas another group received clamped endpoint feedback, which minimizes explicit learning thus isolating implicit adaptation (Brudner et al., 2016; Schween and Hegele, 2017; Morehead et al., 2017).

To measure sensitivity to the error size, we used a modified version of the system-identification approach of varying the direction and magnitude of the rotation pseudorandomly (Scheidt et al., 2001; Fine and Thoroughman, 2006, 2007; Wei and Kording, 2010; Marko et al., 2012; Semrau et al., 2012; Hutter and Taylor, 2018). In the present study, however, the rotation was not changed every single trial but every two trials, as the experiment unfolded as a series of two-throws sequences: participants in both groups (Delayed and Clamped) were instructed to aim straight at the bull’s eye once (the “probe” throw), and then either re-aim or keep aiming straight in the second trial (the “test” throw), respectively. The critical measure of error sensitivity was the degree of change from the probe throw to the test throw, with special interest in how depth information could potentially influence error size interpretation by the explicit and implicit processes.

## Methods

### Participants

40 individuals participated in the experiment (15 females; age between 18 and 35). They were equally divided into two groups of 20 for the motor task, whereas all 40 did the same perceptual task. The sample size for the motor task was determined through power analysis based on a previous study (Hutter and Taylor, 2018) that compared the slope of the error sensitivity function of both explicit and implicit learning. Given an effect size of 1.19, the minimum number of participants required to detect a difference between the two learnings through a two-sample t-test was 19 per group (alpha = .05, power = .95). The participants self-reported normal or corrected-to-normal vision. Handedness was measured with the Edinburgh Handedness Inventory. Participants received either a reimbursement of $ 12/hour or coursework credit for volunteering. The experiment was approved by the Institutional Review Board at Princeton University and each participant gave informed consent prior to the experiment.

### Apparatus and Procedure

Participants sat at a table with their head on a chinrest, which could be adjusted to a comfortable position such that the gaze pointed straight to the horizon. The visual scene was displayed through a head-mounted virtual reality system (HTC Vive) with a resolution of 1440×1600 pixel per eye with a 90 Hz refresh rate. The virtual scenes were modeled in Blender (www.blender.org) and rendered in the viewport of the Unity game engine (unity.com), where they were animated through custom C# routines. The experimenter first measured the participant’s interpupillary distance (IPD) using a digital pupillometer, then helped her or him wear the VR system’s headset, adjusting the spacing between the lenses to match the participant’s IPD. Next, the participant performed a stereovision test using a custom program, followed by a short series of familiarization routines to get acquainted with the virtual environment, and finally the actual experiment began.

### Viewing conditions

The experiment included a perceptual and motor task. In both tasks, the target stimulus was a virtual dartboard and two visual conditions were tested: A No-Depth condition, where the target was presented in isolation against a gray background, and a Depth condition, where the target appeared at the end of a full-scale tavern’s hall (fig. 1A). Although both conditions were viewed in stereovision, they differed substantially in the strength and amount of available depth information. The uniform background of the No-Depth condition did not allow the eyes to resolve binocular disparity across almost the entire visual field. The only exception was represented by the area subtended by the target, whose size was chosen such that, within that region, binocular disparity would be negligible. Under this condition, participants could determine distance by relying primarily on oculomotor information about the orientation of the eyes when fixating the stimulus. The target was presented at either at the near distance of 1 m, where oculomotor signals are reliable, or at the far distance of 7 m, where the eyes are essentially parallel and ocular vergence is completely unreliable (Howard, 2012). The small visual angle subtended by the target also ensured that absolute distance information provided by vertical disparities was negligible (Gillam et al., 1988). In sharp contrast, the environment under the Depth condition was enriched with a wide array of visual features, which ensured a reliable disparity field and also provided plenty of additional monocular cues to depth, including perspective lines, texture gradients, surface occlusions and objects with familiar sizes. In both conditions, the target always appeared in the center of the visual field at eye height. Note in figure 1A that the central portion of the visual scene up to about 30 degrees of visual angle (the wall behind the dartboard) showed the same uniform gray background under both conditions. The size of this area was chosen so to ensure that i) subjects could not have an advantage in the Depth condition by being able to directly compare the target’s location with a neighboring scene element or visual feature; ii) the endpoint feedback during the motor task would always appear against a neutral background (i.e., no depth information) in both conditions (see below).

**Figure 1.**
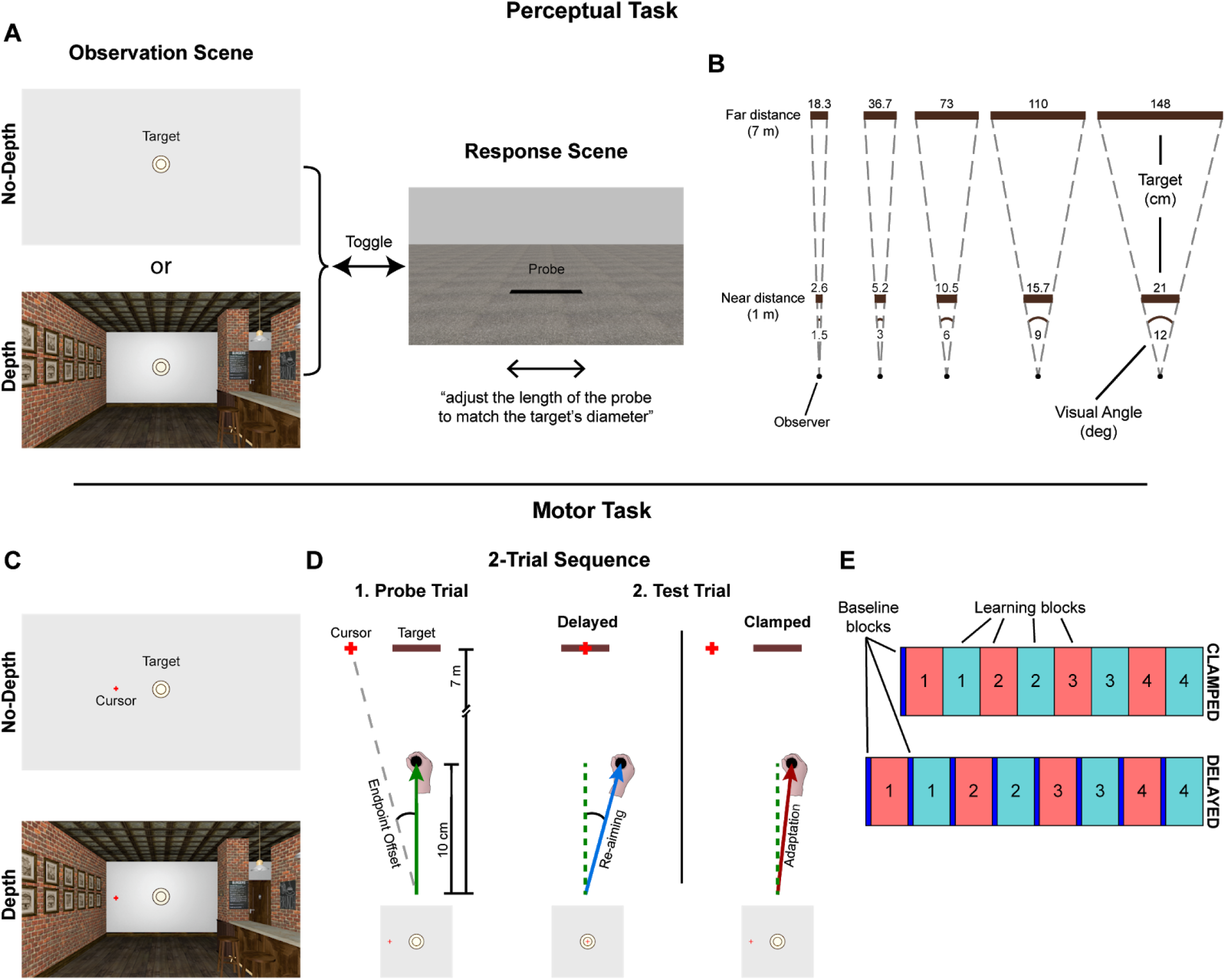
Methods of the study. A) In the perceptual task, participants toggled at will between an Observation scene and a Response scene. During Observation, they saw a dartboard either in isolation (No-Depth condition) or inside a tavern (Depth condition). Only one condition was presented in each trial. In the Response scene, they adjusted the length of a line to estimate the dartboard’s diameter. B) Bird’s eye geometry of the stimulus set in the Observation scene. In each trial, one out of 10 possible target objects was presented (brown rectangles), resulting from the combination of five sizes (in cm in the figure) and two egocentric distances (1 m or 7 m). The size and the distance were calculated such that the target subtended one of five possible visual angles (in deg). Each visual angle was therefore projected from either a near distance or a far distance. C) During the motor task, participants threw virtual darts at the dartboard by making small reaching movements and triggering the appearance of a small cross (Cursor), which provided feedback about the hand’s direction. D) Bird’s eye geometry of the two-trial sequence executed in the motor task. The probe trial (1) came first: All participants were instructed to aim at the target’s center by reaching 10 cm in the forward direction (green arrow), and a red cross appeared at the same depth of the target, showing the hand’s direction plus an angular offset (see text for details). The feedback appeared either one second after movement completion (Delayed group), or immediately following it (Clamped group). The following test trial (2) was different between the groups: The Delayed participants were instructed to re-aim (blue arrow) so to counteract the offset and bring the red cross in the middle of the target; the Clamped participants could not control the endpoint feedback’s position, thus they were told to ignore it and keep aiming straight, which induced involuntary sensorimotor adaptation (red arrow). E) Block design of the two groups of the motor task. The Clamped group (top row) began with a few baseline trials at the beginning of the session, followed by eight consecutive blocks of learning trials, four under each viewing condition (Depth or No-Depth, in different colors). The Delayed group (bottom row) performed baseline trials before each learning block. Both groups rested for 30 seconds between the completion of a learning block and the start of the next one.

### Perceptual task

To determine how depth vision influences the perception of size, participants performed a perceptual task in which they reported the apparent size of the target (the dartboard) while being immersed in either a depth-enriched or a depth-impoverished virtual environment (fig. 1A). Each trial started with the “Observation” scene, that is, with the target visible under one of the viewing conditions (Depth or No-Depth). The target was seen either at 1 meter (near distance) or at 7 meters (far distance) and subtended one of five visual angles: 1.5, 3, 6, 9 or 12 degrees, corresponding approximately to a diameter of 2.6, 5.2, 10.5, 15.7 and 21 cm at the near distance, and of 18.3, 36.7, 73, 110 and 148 cm at the far distance (fig. 1B). Through a keypress, participants could toggle between this scene and a second scene (“Response”). The Response scene showed a textured floor with a probe laying on its surface 4 meters away from the observer. The highly detailed surface of the ground in the Response scene was designed to provide participants with a reliable reference when reporting their estimates of distance and size from the Observation scene. The probe’s distance was chosen to be intermediate between the target’s near and far distances. The probe was a black strip orthogonal to the line of sight. While in the Response scene, the participants used the keyboard to adjust the length of the probe until it matched the perceived diameter of the target in the Observation scene. Participants were instructed to imagine that the target was illuminated by perfectly vertical sunlight, and that the probe was to represent the resulting shadow being cast on the floor. Using the keyboard, participants could switch back and forth between Observation and Response scenes as many times as they needed. Following a within-subject design, the task included five presentations of each of the five target’s sizes at each distance in each condition, for a total of 100 trials.

### Motor task

To determine if depth vision affects how visual errors are encoded by the motor system, participants performed a visuomotor adaptation task, in which they were instructed to throw a virtual dart at the target’s bull’s eye while being immersed in either a depth-enriched or a depth-impoverished virtual environment. They did so by making short reaching movements to slide one of the HTC Vive controllers on a tabletop, starting from near the chest and moving in forwards direction. The controller was installed on a custom 3D-printed support whose basis was covered in felt and glided on a glass surface, to minimize friction and auditory cues. The experimenter first calibrated the start position, then the participant donned the VR headset, rested their head on the chinrest and practiced reaching out and returning to start position until the movement became smooth and natural. Akin to throwing a real dart, participants made short rapid 10 cm reaching movements with the hand as soon as the dartboard appeared (median movement duration = 464 msec). When the hand was 10 cm away from the start position along the depth dimension, a 1.5 degrees wide cross appeared on the screen (fig. 1C). The cross was located at the same distance of the target and at the same height of the target’s center (at eye height), while its lateral position provided endpoint feedback about the movement direction (fig. 1D). No online feedback was given. The goal of the experiment was to study the influence of pictorial depth information, such as perspective and texture, during motor learning. Since binocular disparity tends to override these cues altogether, and it is most reliable near the body (see also below the results from the perceptual task), in the motor task the target was always seen at the far distance (7 m) subtending a visual angle of 6 degrees (diameter of 73 cm).

#### i. Experimental groups

While the perceptual task was identical for all participants, in the motor task half of the participants were assigned to a “Clamped” group, presumed to be largely implicit, and the other half to a “Delayed” group, presumed to be mostly explicit. Both groups performed eight blocks of trials back-to-back with a short 30 seconds break between each block. In a within-subject design, four blocks tested reaching under the Depth condition, and four under the No-Depth condition. For the Delayed group, each block started with a short sequence of “baseline” trials followed by a long sequence of “learning” trials. Clamped participants performed a series of baseline trials only at the beginning of the first block.

#### ii. Baseline trials

During a baseline trial, the target appeared under one of the two viewing conditions and subjects threw one virtual dart at its bull’s eye, receiving veridical endpoint feedback about their movement’s accuracy. For the Clamped group, feedback was displayed immediately after the hand crossed a distance of 10 cm. For the Delayed group, endpoint feedback was presented 1 second after the hand crossed 10 cm. For both groups, the feedback then remained visible for 1 second, after which the scene was cleared, and the subject could return to the start position to begin the next trial. A V-shaped metallic frame was installed at the start position to guide participants to the same initial location when returning.

#### iii. Learning trials

Each learning trial consisted of two throws. In the first throw, the “probe”, participants of both groups were instructed to aim at the center of the bull’s eye as accurately as possible, but the endpoint feedback was offset laterally by the computer by a given amount of degrees of visual angle. There were eleven offsets: 0, ±1.5, ±3, ±6, ±9 or ±12 degrees. For the Clamped group, the position of the endpoint feedback was independent of the actual hand direction. For example, with a zero degree offset the endpoint feedback appeared exactly in the center of the target (i.e., no visual error), regardless of where the subject actually reached with the hand. Similarly, when the offset was -9 degrees, the endpoint feedback appeared 9 degrees to the left of the target’s center (i.e., bullseye), regardless of the participant’s hand direction. Subjects in this group were instructed to ignore the endpoint feedback and always aim at the target’s center.

For the Delayed group, the endpoint feedback’s position resulted from shifting the true hand direction by an amount corresponding to the offset size. For example, a zero degree offset meant that the endpoint feedback was veridical (the actual direction of the reaching plus zero offset), and a +9 degrees offset meant that the true hand direction was shifted 9 degrees to the right. Subjects in this group were instructed to aim straight at the target’s bullseye and pay attention to the endpoint feedback.

In the second throw of the learning trial, the “test”, the endpoint feedback’s behavior was the same as in the probe (fixed for the Clamped participants and shifted for the Delayed participants), but the instructions differed between groups. The Clamped group was instructed to aim again at the target’s bullseye and keep ignoring the endpoint feedback, while the Delayed group was instructed to counteract the offset observed in the probe throw, such as to bring the (shifted) endpoint feedback as close as possible to hitting the target’s center. Like in the baseline trials, endpoint feedback on learning trials appeared either instantaneously (Clamped) or with a 1 second delay (Delayed) and remained visible for 1 second. To ensure that the participants kept aiming at the target’s center, every trial (i.e., every sequence of a probe and a test throw) the target’s horizontal position was extracted from a uniform distribution within the interval [-1.5, 1.5] degrees (the computer-controlled endpoint feedback of the Clamped group was also shifted accordingly). This manipulation also prevented that participants could exploit the same absolute target location to estimate the position of the endpoint feedback.

For the Clamped group, the go-signal was a beep sound, and the appearance of the endpoint feedback was accompanied by a brief thud. Once a block of learning trials was completed, the program cleared the visual scene and the subject rested 30 seconds while listening to background music before starting a new block. The go-signal for the Delayed group was a recorded voice saying “shoot” (baseline trials), “probe” (first throw of the learning trial) or “test” (second throw of the learning trial). The appearance of the endpoint feedback was accompanied by a tone which increased in pitch as the feedback hit closer to the target’s center. Furthermore, subjects also scored between 0 to 100 points each trial depending on how accurate their reaching was. During the 30 seconds break following the end of a given block, Delayed participants were shown a chart where they could compare their ranking against that of other nine virtual opponents up to that point and were encouraged to “win” the game.

### Experimental design

Participants in both groups performed 264 learning trials. Each learning trial comprised two reaching movements (a probe trial followed by a test trial) – 2 viewing conditions x 11 offsets x 12 repetitions (amounting to 528 throws) – distributed across eight blocks (four blocks per visual condition).

The order of the viewing conditions (Depth and No-Depth) over the course of the eight blocks was counterbalanced between subjects, as well as the order of the tasks (perceptual task first versus motor task first). The Delayed group performed an additional 14 baseline trials at the beginning of each block to emphasize re-aiming in the subsequent learning block. For the opposite reason, namely to avoid canceling implicit adaptation, the Clamped group did only 10 baseline trials at the beginning of the experiment (fig. 1E).

### Data Analysis

The responses of the perceptual task and the kinematic data were analyzed in R (R Core Team, 2020). For the perceptual task, the main dependent variable was the adjusted size of the probe object in the Response scene. An exploratory data analysis was first performed on a subject-by-subject basis for the detection of possible outliers, defined as those adjustments which exceeded 1.5 times the IQR of the total distribution of the responses for that individual. These observations were excluded, amounting to 0.9% of the entire data set.

For the motor task, the primary dependent variable was the lateral position of the hand when it reached 10 cm in depth from the start position, which the program recorded in real time and used to project the endpoint feedback. From that value, the instantaneous hand direction was calculated as the angular deviation from the forward direction (positive values for clockwise deviations). In addition, the 2D positional data of the controller (x and z coordinates of the reaching movement) was also examined. First it was checked for missing frames, on a subject-by-subject and trial-by-trial basis. A Savitzky-Golay filter of fourth order was then applied to the trajectory data and used to compute velocity and acceleration. The onset of the reaching movement was chosen to be the first frame of the longest sequence of frames in which the velocity’s z component (the velocity in the forward direction) was continuously greater than 2 cm/s. Within the same sequence, the end of the movement was marked by the first frame in which the z velocity dropped below 2 cm/s. From the processed trajectory data, we extracted a number of dependent variables, including velocity and acceleration maxima and their times of occurrence.

## Results

### Perceptual Task

The perceptual task measured how size perception is influenced by the amount of visual depth cues present in the environment. In it, participants adjusted the length of a line to match the perceived diameter of a circular target object (a dartboard) which was seen either inside a highly detailed room (Depth condition) or in an empty space (No-Depth condition). Every individual did the same perceptual task regardless of the motor task group they were assigned to.

On average, participants increased the length of the probe in a semi-linear fashion as the visual angle subtended by the target increased, as shown in figure 2. The five target’s angular sizes were easily distinguishable from one another, as confirmed by the Holm-corrected post-hoc comparison of all pairwise differences between the adjusted sizes. Furthermore, in agreement with the psychophysics literature (Oyama, 1974; Ross and Plug, 1994), the just-noticeable-difference decreased for larger stimuli, namely the same size interval appeared smaller between two large stimuli than between two small stimuli. This led to a gradual departure from linearity in the responses as a function of the visual angle subtended by the target (fig. 2). The critical test here was to see whether the different depth contexts would affect this size estimation and how. The angular size of the probe’s adjusted length, matching the perceived angular size of the target, was submitted to an omnibus univariate 5×2×2 ANOVA, crossing three factors: the target’s simulated size, the viewing distance and the viewing condition. All three main effects were significant (target size: F_(4,156)_ = 558.96, p < .001, 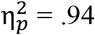; viewing distance: F_(1,39)_ = 252.17, p < .001, 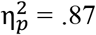; visual condition: F_(1,39)_ = 39.64, p < .001, 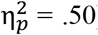), as well as all the interactions (viewing distance X target size: F, = 30.32, p < .001, 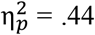; viewing distance X visual condition: F_(1,39)_ = 56.62, p < .001, 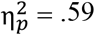; target size X visual condition: F_(4,156)_ = 13.10, p < .001, 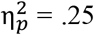; viewing distance X target size X visual condition: F_(4,156)_ = 7.72, p < .001, 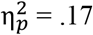).

**Figure 2.**
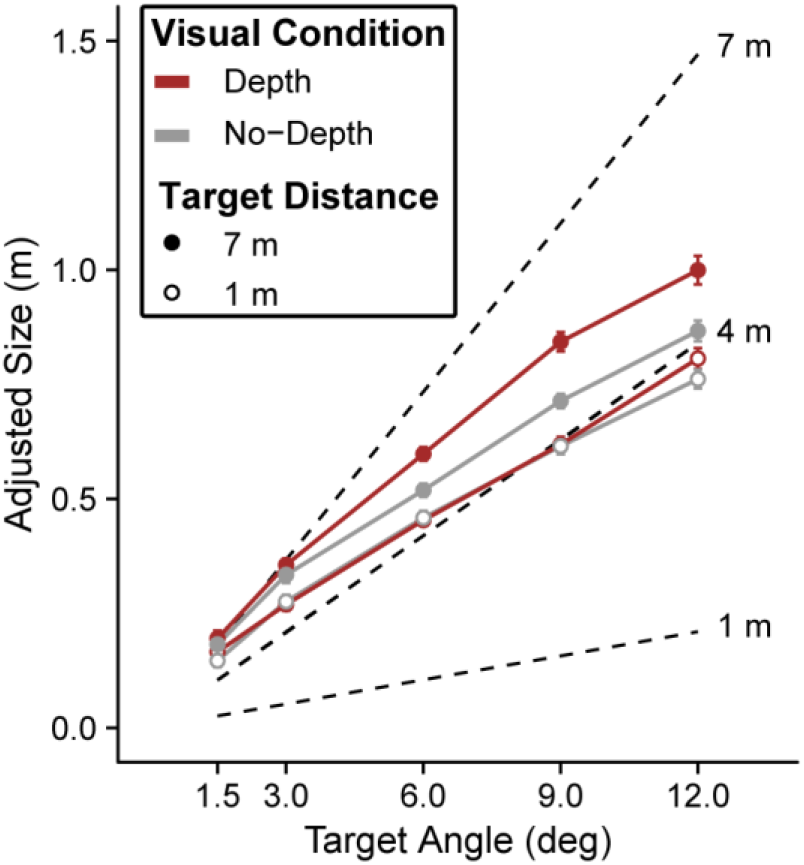
Adjusted size of the probe in the perceptual task as function of the stimuli’s visual angle. Adjustments of the far (7 m) and the near (1 m) stimuli are represented by filled circles and open circles, respectively. Two dashed lines represent the target’s simulated sizes at the two distances. A third dashed line indicates the target’s projected sizes at the probe distance of 4 m (at which the adjustments were made). Red data points and solid lines identify the responses under the Depth condition, while the responses under the No-Depth condition are displayed in gray. Error bars represent the within-subject 95%CI.

The main effect of the viewing condition showed that the size estimates were significantly greater when the target was seen under the Depth condition than in isolation (F_(1,39)_ = 39.64, p < .001). Figure 2 shows that this effect was primarily driven by the interaction with the fixation distance, as it was significant at 7 m (F_(1,39)_ = 71.14, p < .001; solid dots), but not at 1 m (F_(1,39)_ = 2.47, p = .12; open circles). This was expected, since at 7 meters the pictorial cues (perspective, texture, relative size, et cetera) dominated size perception over binocular disparity, which instead decays rapidly with the square of the distance (Howard and Rogers, 2002). As a result, when the dartboard was seen at the far distance in the richly detailed scene of the Depth condition, it appeared compellingly larger (because seemingly more distant) than when seen at the same distance against the flat background of the No-Depth condition. For this reason, 7 m was also the distance selected for the motor task.

The reverse side of the interaction between depth cues and fixation distance was that, under the Depth condition, the difference between the perceived size of the far objects and that of the near objects was greater relative to No-Depth (figure 2, solid dots versus open circles for each color). The abundance of depth cues in the former condition allowed participants to distinguish between the two fixation distances more clearly, resulting in more separate size estimates at near and at far distances. This pattern is in accordance with a known compression/expansion mechanism affecting the perceived visual space which depends on the amount of depth cues available in the environment (Campagnoli et al., 2017; see discussion).

In summary, size estimates were systematically greater when the target was seen in the tavern scene than when seen against a uniform background, and this effect was maximal at the farthest fixation distance as anticipated.

### Motor Task

In the motor task, participants performed a visuomotor rotation task aiming at a target seen at eye height from a distance of 7 m. To focus selectively on explicit and implicit components of motor learning, they were divided between a Delayed group and a Clamped group, respectively. To study if and how the two processes are affected by changes in the depth information in the environment, both groups did the motor task under a Depth and a No-Depth visual condition. Throughout this task, all participants performed three types of trials (baseline, probe and test). The direction of their reaching movements, which we will refer to as hand angle, was measured as the angle between the instantaneous direction of the hand when it had traveled 10 cm away from the body and the forward direction.

All participants aimed straight during the baseline trials, namely with an average hand angle not different from zero (Delayed: -.004 ± .007 deg, t_(19)_ = -.65, p = .52; Clamped: .03 ± .04 deg, t_(19)_ = .73, p = .47). In the learning blocks, the average hand angle of the probe trial was also zero in both groups, as expected (Delayed: .06 ± .1 deg, t_(19)_ = .64, p = .53; Clamped: -.14 ± .13 deg, t_(19)_ = -1.12, p = .28). While both groups completed these movements in the same amount of time (Delayed: 510 ± 40 ms; Clamped: 460 ± 40 ms; t_(38)_ = .83, p = .41), Clamped group exhibited shorter RTs than Delayed group (Clamped: 364 ± 44 ms; Delayed: 642 ± 61 ms; t_(38)_ = 3.69, p < .01), as the latter group actively tried to counter the visual rotation. Finally, in the test trials, the hand angle was again on average zero (Delayed: .18 ± .16 deg, t_(19)_ = 1.13, p = .27; Clamped: -.2 ± .14 deg, t_(19)_ = -1.4, p = .18) since the distribution of the offsets was symmetrical around the straight direction. Moreover, since the test trials alternated with the probe trials during the learning phase of the experiment, their timing followed the same pattern (same movement time in both groups, shorted RTs for the Clamped group).

To evaluate the effect of the experimental design on motor learning, a 2×6×2 mixed ANOVA was performed on the hand angle of the test trials, crossing one between-subjects factor (group) with two within-subject factors (offset size and visual condition). All three main effects were significant (group: F_(1,38)_ = 444.95, p < .01, 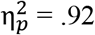; offset size: F(5,190) = 408.42, p < .01, 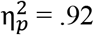; visual conditions: F_(1,38)_ = 4.56, p = .04, 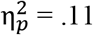), as well as the interactions between group and offset size (F_(5,190)_ = 360.35, p < .01, 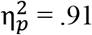) and between group and visual condition (F(1,38) = 13.96, p < .01, 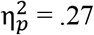).

As the main effect of the offset size shows, both groups exhibited learning. The main variable we used to quantify learning throughout the whole study was the individual sensitivity to the error size, computed through a subject-by-subject linear regression of the hand angle as a function of the offset. This analysis returned an average slope of .96 (t_(19)_ = 23.98, p < .01) for the Delayed group, and an average slope of .04 (t_(19)_ = 3.88, p < .01) for the Clamped group (fig. 3). Though decidedly smaller than the effect size of the explicit learning, the significant scaling with the error size exhibited by the Clamped group confirms previous evidence that the motor system can adapt to perturbations of the visuomotor mapping that vary as frequently as every other trial (Fine and Thoroughman, 2006, 2007; Wei and Kording, 2009; Marko et al 2012; Semrau et al., 2012; Hutter and Taylor, 2018). Moreover, the Clamped group showed learning despite that certain features of the task could have potentially reduced its magnitude, like receiving endpoint rather than continuous feedback (Hinder et al 2008; Taylor et al., 2014), and the fact that the experimental setup required converting the error estimate from the frontoparallel plane (that of the visual input) to the transversal plane (that of the hand).

**Figure 3.**
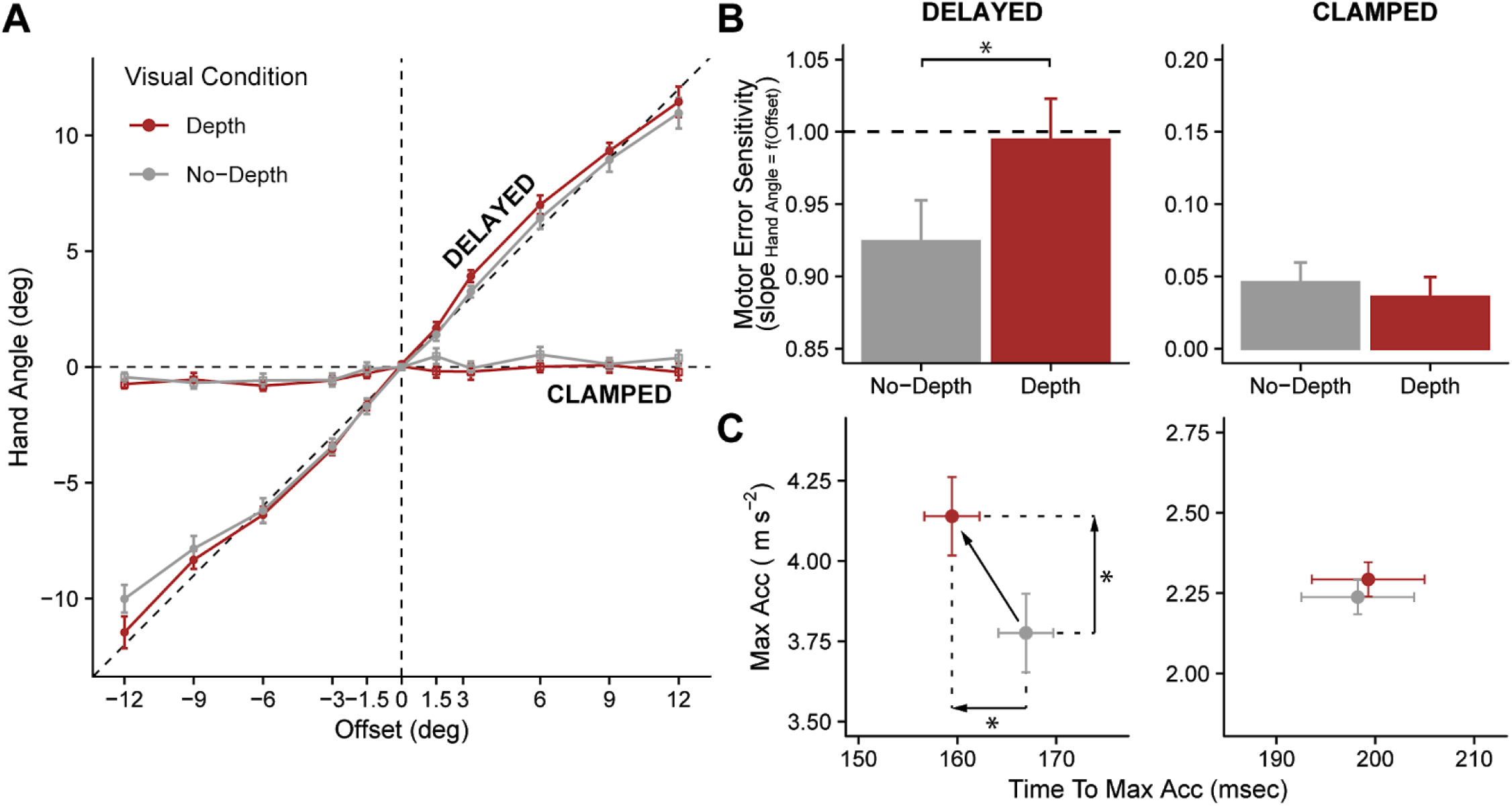
A) Hand angle of the test trials as a function of the offset presented in the probe throw, for the Delayed group (circles) and the Clamped group (squares). Error bars represent the within-subject SE of the mean. Across all panels, the same color code is used to indicate the visual conditions (Depth in red and No-Depth in gray). Panels B and C show additional statistics about the performance of the two groups – left, Delayed; right, Clamped. The aspect ratio is the same across each row of panels. B) Significant increase in error sensitivity under the Depth condition relative to No-Depth (for Delayed but not for Clamped), expressed through the slope of the linear regression relating the hand angle to the offset size; C) Significant anticipation (x axis) and increase in magnitude (y axis) of the peak acceleration under Depth relative to No-Depth (for Delayed but not for Clamped), compatible with a movement aimed at a farther target in the former condition.

The most critical finding of this task, the main effect of the visual conditions, was caused by the Depth condition yielding larger compensations than the No-Depth condition (fig. 3A). This behavior mirrored the bias in the estimation of the target’s size in the perceptual task: relative to the nondescript environment, participants showed greater compensations when throwing virtual darts in the tavern scene.

More precisely, this main effect was driven by its interaction with the group factor, as the visual condition had a significant effect on the performance of the Delayed group (F_(1,38)_ = 17.24, p < .01) although not for the Clamped group (F_(1,38)_ = 1.28, p = .26). This difference between the responses of the two groups in the test trials can also be viewed in terms of error sensitivity, calculated as the slope of the linear regression between the hand angle and the offset size (Fig. 3B). For the Delayed participants, switching to the tavern environment caused sensitivity to significantly increase by .08 (t_(19)_ = 3.5, p < .01) relative to the nondescript environment, reaching a slope value of .99. For comparison, no significant change could be found between conditions for the Clamped group (t_(19)_ = -.94, p = .82).

While we cannot rule out that the small magnitude of the implicit adaptation could have prevented a depth-related effect to be detectable, the hand angle data could also hint to a genuine dissociation between explicit and implicit processes with respect to how depth information is integrated for the encoding of error signals. The analysis of two other behavioral markers provided partial evidence in favor of the latter possibility, suggesting that indeed depth vision selectively affected the movement plan of the Delayed group only.

First, participants in the Delayed group exhibited faster movements when throwing darts under the Depth condition than under the No-Depth condition (fig. 3C), consistent with them aiming at a target that appeared more distant in the former case, and in agreement with the results of the perceptual task. The peak acceleration of the test throw was greater (t_(19)_ = 2.1, p = .02) and also occurred earlier in time (t_(19)_ = -2.21, p = .02). In contrast, the Clamped group showed no change between conditions neither in the peak acceleration’s magnitude (t_(19)_ = .73, p = .24) nor in its time of occurrence (t_(19)_ = -.38, p = .35).

Second, based on the previous results we hypothesized that the reaction times would be greater under the Depth condition than under No-Depth, due to increased mental rotation required to compute a perceptually bigger re-aiming angle (McDougle and Taylor, 2019). To test this prediction, we regressed the reaction times on the offset size (absolute value) for each subject, using the resulting slope as a measure of mental rotation. To calculate the net mental rotation in the test trial, we subtracted the fit of the probe trial data from that of the test trial. We calculated two such values for each subject, one for each visual condition. This dataset was submitted to a one-way ANOVA with the visual condition as the within-subject predictor, finding a marginal effect of the visual condition in the Delayed group (F_(1,19)_ = 4.48, p = .048, 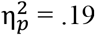), and no effect in the Clamped group (F_(1,19)_ = .10, p = .75, 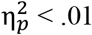).

### Coupling between perception and motor learning

To assess the degree to which motor learning relies on depth information, we took a closer look at the relationship between the responses in the motor task and the perceptual judgments. Given the small effect size of the hand angle results of the Clamped group, we excluded that data from this analysis. The possible relationships between depth information and the different components of motor learning are addresses in the discussion.

An analysis comparing the motor and the perceptual data showed that the extent to which the Delayed participants adjusted their motor performance under the Depth and No-Depth visual conditions closely reflected their own perceptual judgments under the same conditions. In a series of post-hoc tests, we broke down the Delayed motor data into each learning block and looked at its evolution over time. As per the experimental design, the Depth and No-Depth environments were each presented in four successive learning blocks across the motor task (fig. 1E), enabling us to compare how the error size sensitivity varied on a block-by-block basis in each visual condition.

The average .08 relative increase in slope under the Depth condition relative to No-Depth was not constant, but it grew almost linearly over the course of the four learning blocks. More precisely, this gradual change in sensitivity was not caused by an absolute improvement of the performance under the Depth condition, but rather to its deterioration under the No-Depth condition (fig. 4A). We fitted the error size sensitivity as function of the number of blocks for each condition, and found that it was accurate (intercept = 1.02, t_difference from 1(19)_ = .44, p = .66) and constant (slope = -.003, t_(19)_ = -.14, p = .89) under Depth, whereas it decreased significantly under No-Depth (slope = -.04, t_(19)_ = -2.66, p < .01).

**Figure 4.**
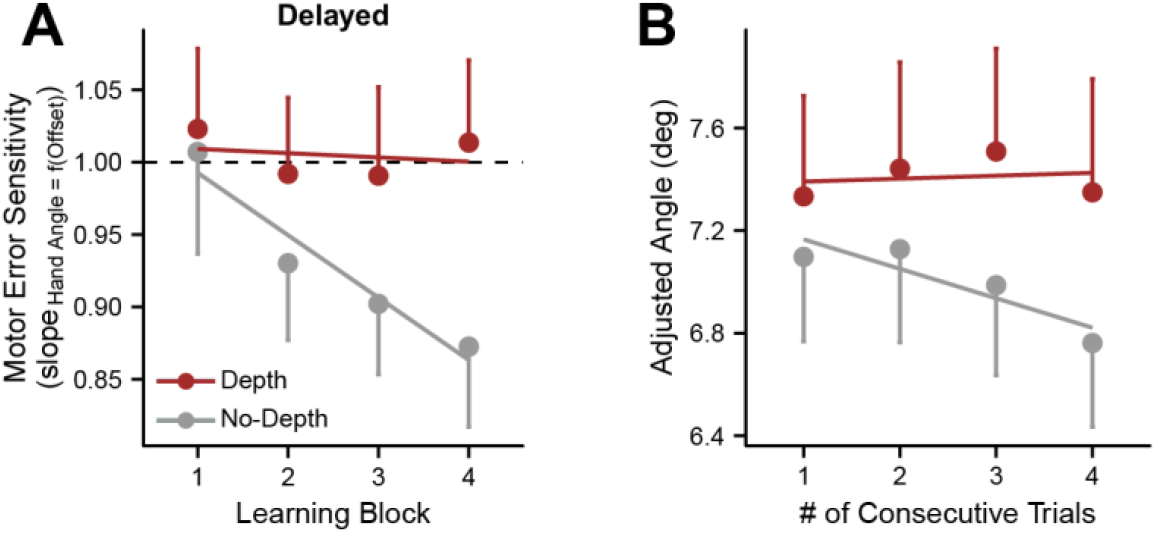
A) Motor task: Block-by-block evolution of the motor error sensitivity for each visual condition (Delayed group). B) Perceptual task: Average estimate of the target size as function of the number of trials consecutively under the same visual conditions. All error bars show the within-subject SEM.

If the motor error sensitivity and the perceptual error sensitivity stemmed from the same process, we hypothesized that an analogous pattern should be found also in the perceptual results. To this end, we analyzed the temporal evolution of the same participants’ judgments of size. Since the perceptual task consisted of a unique block, we looked at how the responses changed on a trial-by-trial basis. For each subject we split the dataset into sequences of trials in which the same visual condition was presented consecutively due to randomization. We were able to obtain a full sample size for strings up to a length of 4 trials. The comparison between this dataset and the motor dataset uncovered two notable results.

First, we found yet another group-level similarity between the motor and the perceptual data. Specifically, longer sequences of size judgments made under the same visual condition correlated with a stronger effect of depth information and vice versa. To compute this effect, we fitted a subject-by-subject linear model of the average adjustments as a function of the number of consecutive trials under a given condition. We found a significant interaction between the trials sequence’s length and the visual condition, such that the No-Depth adjustments became significantly smaller for longer sequences of consecutive trials (t_(39)_ = -1.88, p = .03), whereas the Depth adjustments remained constant (t_(39)_ =.39, p = .70) (fig. 4B), attributable to size adaptation (see Discussion). Though these results are only indirectly confirmatory, as they show a correlation between a block-by-block effect with a trial-by-trial effect, they nonetheless reveal the signature of the same mechanism affecting, on a different scale, both the perceptual and the motor results.

Second, and more critical, evidence of a perception-action coupling was found at the individual level. As the top row of figure 5A shows, the error size sensitivity under the No-Depth condition showed a maximum, nearly significant drop in the second block (t_(19)_ = -1.47, p = .078), which is evidence of a sharp underestimation of the target’s distance relative to the first block. This decrease then abated in the remaining third and fourth blocks (gray data). For comparison, under the Depth condition (red data), the transition from the first block to the second block was accompanied by a much smaller drop in error sensitivity (p = .27).

**Figure 5.**
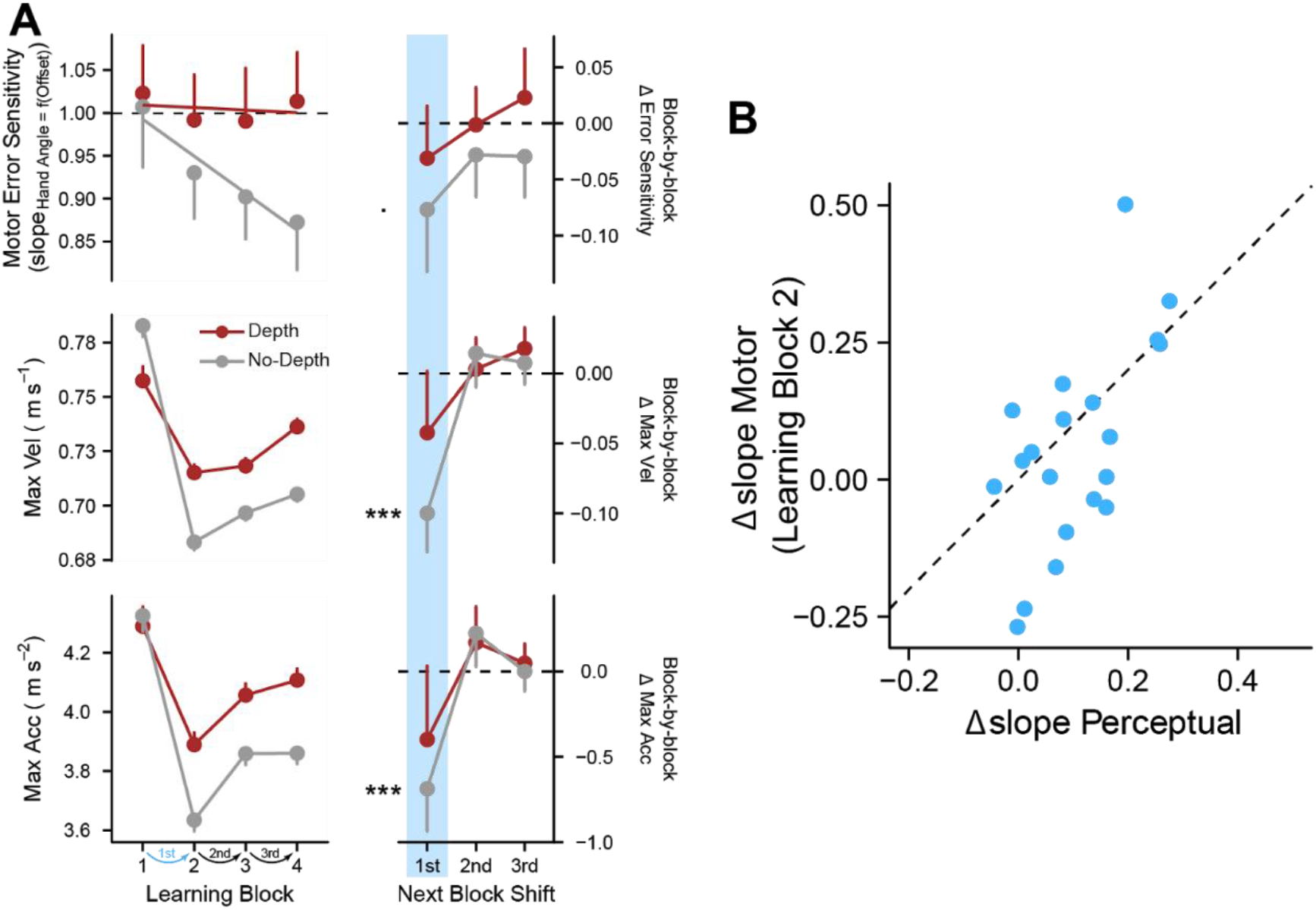
A) Group-level effects of depth information on motor learning (Delayed participants). Left column, evolution of three behavioral indexes (top row, motor error sensitivity – same as in figure 4A; middle, max velocity; bottom, max acceleration) over the course of the learning blocks of each visual condition (Depth in red, No-Depth in gray). Right column, block-by-block difference of each variable (change relative to the previous block): most of the motor learning was determined after shifting to the second block. Error bars show the within-subject SEM. B) Individual-level effect of depth information on the perception-action coupling. Comparison between the effect of the visual conditions on the error sensitivity (Δslope) in the motor task (y axis) and in the perceptual task (x axis) during learning block #2. Each point represents an individual score.

Although at first blush this too could be viewed as completely spurious, but a careful observation of the movement kinematics revealed the presence of the exact same pattern (fig. 5A, center and bottom rows). Under the No-Depth condition, both the hand’s maximum velocity and maximum acceleration underwent a dramatic reduction after the first transition from the first block to the second block (max velocity: t_(19)_ = -3.33, p < .01; max acceleration: t_(19)_ = -2.60, p < .01), consistent with the reduction of the target’s estimated distance suggested by the analysis of the error sensitivity. Moreover, this means that the error was lawfully underestimated in turn, since the offset size scaled with the viewing distance. In the remaining blocks of the motor task, the kinematic profile of the hand then remained relatively stable, again mirroring the error sensitivity: both maxima showed no significant reduction under the Depth condition (max velocity: p = .20; max acceleration: p = .20).

Note in the graphs in figure 5A that, although there was a general reduction of all behavioral indices in the second block (all trended downwards), likely due to habituation to the visual scene, this very reduction was always systematically greater in the No-Depth condition than the Depth condition.

All these observations suggested that the first shift between blocks under different visual conditions was the most critical, in terms of when the depth cues had their most visible impact on error sensitivity. To verify this conclusion, we repeated the previous correlation analysis between the motor and the perceptual shift in error sensitivity (Δslope) with one important modification: rather than taking the subject-by-subject motor data averaged across all four learning blocks, we subset each learning block and compared it with the same individuals’ perceptual results. This resulted in four correlation statistics (Pearson’s r), comparing the same Δslope_perceptual_ with the Δslope_motor_ from the first, second, third and fourth motor learning blocks. Consistent with the behavioral observations, we found that the perceptual data showed a selective and strong positive correlation with the motor data from the second block (r_block2_ = .59, p = .02) – that is, after the first shift between visual conditions but not after the following shifts (p_block1_ = .25, p_block3_ = 1; p_block4_ = 1; all p-values were Bonferroni-corrected).

This analysis outlined the importance of the first transition from one condition block to the other, which appeared to mark a crucial moment during explicit motor learning. In the second learning block, each Delayed participant recalibrated their reaching movements according to their own perceptual rescaling of the scene. This is visible in the scatterplot of figure 5B: if, for example, a participant estimated the target’s size in approximately the same way under either visual condition, showing a weak effect of the depth information on perception – that is, a low Δslope_perceptual_ – they also showed a weak effect of the depth information on action – a low Δslope_motor_. Vice versa, a participant who showed a clear improvement in the estimation of the target’s size under the Depth condition relative to No-Depth – resulting in a high Δslope_perceptual_ – also scaled their re-aims more accurately under the same visual condition (high Δslope_motor_).

Overall, this final analysis of the temporal evolution of the results uncovered compelling parallelisms between perceptual and motor data. Although these similarities must be taken with caution due to the post-hoc nature of the analysis, they are remarkably consistent with the results of the contrasts planned *a priori*. In conclusion, while more future exploration on this issue is necessary, the present data strongly supports a direct connection between the perceptual analysis of the visual scene and the behavior exposed by explicit motor learning.

## Discussion

Behavioral and neuroanatomical research on motor adaptation have uncovered evidence of both explicit and implicit error-minimization mechanisms, which drive the human ability to adapt to changes in the environment. Both processes are known to react to error signals differently from one another (Taylor et al., 2014; McDougle et al., 2015; Bond and Taylor, 2015; McDougle and Taylor, 2019). Regardless of these behavioral differences, any incoming source of error information requires translating a proximal sensory stimulation into a representation that is useful to update the motor output.

Theoretically, the translation step poses a problem, because it happens in the perceptual domain and perception does not always represent reality in a veridical fashion (Johnston, 1991; Todd et al., 1995; Bingham et al., 2000; Todd and Norman, 2003). In practice, the effects of potentially inaccurate perception-action coupling have not been a concern for most of the previous investigations, which have probed mostly planar movements with feedback seen on a flat surface or a frontoparallel setup. However, in the natural environment motor actions are embedded in a 3D space, and they frequently consist of movements aimed at multiple distances where even small perceptual changes can lead to major mistakes. Here we sought to investigate the role of depth information in motor learning using virtual reality to render a 3D version of a classic visuomotor rotation task in depth.

Participants performed outward reaching movements to “throw” virtual darts at a dartboard (the target) located at a simulated distance of 7 meters. They did so under two visual conditions: A Depth condition, where the environment was enriched with retinal and extraretinal depth cues, and a No-Depth condition, where the environment offered no reliable distance cues except for oculomotor convergence. We crossed these visual conditions with two feedback conditions, delayed feedback and “clamped” feedback, which have been shown to emphasize learning by explicit and implicit processes, respectively (Kitazawa et al., 1995; Honda et al., 2012; Brudner et al., 2016; Schween and Hegele, 2017; Morehead et al., 2017; Kim et al., 2018, 2019; Poh and Taylor, 2019).

The learning task consisted of two reaching movements in sequence, a probe trial and a test trial. In the first trial (probe) participants aimed straight at the target’s bullseye and witnessed the perturbation, according to their feedback condition. In the second trial (test), the Delayed group was allowed to re-aim in order to compensate for the offset, whereas the Clamped group was instructed to ignore the feedback and keep aiming straight. Finally, to assess the effect of the visual conditions on size estimation, all participants did a perceptual task where they judged the size of the target under the Depth and the No-Depth conditions.

Both groups of participants exhibited sensitivity to the error size, evidence of learning, although to different extents. Delay participants made ample and mostly accurate corrections, whereas the Clamped participants exhibited a small subconscious compensatory bias proportional to the size of the visual offset. The different error sensitivity between groups was expected, since explicit re-aiming and implicit adaptation have been previously shown to lead to separate forms of learning, each with specific characteristics (Mazzoni and Krakauer, 2006; Taylor and Ivry, 2011; Morehead et al., 2017; Butcher and Taylor, 2018).

More importantly, the analysis of the performance in the test trials highlighted two main results. First, Delayed participants showed a systematic modulation of their error sensitivity driven by the depth condition, such that their re-aiming direction was smaller under No-Depth relative to Depth and correlated with a reduction in the movement’s speed and acceleration. Albeit being relatively small, the decrease in error sensitivity was present for both rightward and leftward errors of the cursor, thus somewhat acting as an internal replication, and we expect that the same increase would be more pronounced with stronger depth cues and at larger distances. This effect of the depth cues suggests that, during explicit motor learning, the error size (i.e., the amount of lateral shift of the endpoint from the target’s center) was estimated as a lateral offset. That is, by scaling its retinal projection by some estimate of the fixation distance to return a length estimate. This would explain why the hand direction was susceptible to changes in the context: as per the perceptual results, perceived size was smaller in the No-Depth environment relative to Depth, yielding less re-aiming, because the overwhelming lack of depth cues caused the flattening of the perceptual space (Foley, 1980) and made the objects in front of the observer look closer and thus smaller. The opposite happened when the scene was abundant with depth cues. The comparison between the motor responses and the perceptual reports provides converging evidence of a coupling mechanism governing distance estimation for both explicit learning and perception. We discuss these findings in the section below.

Second, in contrast to the Delay participants, the error sensitivity function of the Clamped group, as well as their movement kinematics, appeared to be independent of the visual conditions. This selective effect of depth vision on the explicit but not on the implicit learning is a finding that bears further investigation. In this study, important behavioral markers related to the target’s perceived distance seemed to indicate that depth information had no effect on sensorimotor adaptation. Indeed, contrary to the Delayed group, the Clamped participants exhibited no modulation neither of the peak acceleration (related to the target’s perceived distance) nor of the reaction times (related to the error’s estimated size) between visual conditions. However, there are also valid reasons to believe that this null effect may just be an absence of evidence rather than evidence of absence. The effects of the depth cues might have been too small be detected, given how flat the error sensitivity function was for the Clamped group. Or maybe the depth cues were concentrated too much in the periphery of the visual field relative to the area where the clamped feedback appeared. At this stage, there are merits to both accounts. On the one hand, a common mechanism of depth analysis shared by both explicit and implicit learnings would be in principle a parsimonious, thus preferable, account. On the other hand, there is potential support in the literature to the hypothesis that explicit and implicit processing of depth information could feed from different encodings of the depth signals. We explore this evidence below.

### Evidence of coupling between explicit motor learning and perception

The correlation between the error sensitivity of the Delay participants and their size judgments (fig. 5B) is a compelling and straightforward evidence of a relationship between explicit learning and perception, but it is not the only commonality that was found. A second example is the relationship with time shown in figure 4. As previously described, the No-Depth scene was perceptually highly ambiguous: aside from the oculomotor signal, which is rather noisy, the available retinal information allowed for infinite estimates of the target’s distance and size, and by extension, of the offset size. Due to the lack of reliable depth information, the visual system was biased towards pulling the target closer to the body than what was actually simulated, according to the specific distance tendency (Gogel, 1969), leading to size underestimation in adherence with Emmert’s law. In addition, the repeated exposure to the same stimulus gradually induced size adaptation, causing the progressive decrease in apparent size (Köhler and Wallach, 1944; Sutherland, 1961). For the opposite reason, the 3D interpretation of the Depth scene was highly constrained by a rich array of depth cues, thus more perceptually stable. This phenomenon is visible in figure 4A, and in figure 4B we found the signature of the same effect in the perceptual data.

Size misjudgments are not the only bias that can emerge from inaccurate estimation of the fixation distance. Prior to retinal information, the most readily available cue-to-distance is the oculomotor sensation of rotating the eyes to gaze at a new location. The No-Depth condition was designed to measure the contribution of this extraretinal signal essentially in isolation. The results of the perceptual task show that under No-Depth, the adjusted size was statistically greater at far than at near (F_(1,39)_ = 121.26, p < .001), indicating that participants indeed scaled the target’s retinal projection based on detectable changes in vergence. This was expected, since the near and the far distances required two noticeably different eye positions. However, a closer look at the data revealed that the diameter of the near objects had been largely overestimated (t_(39)_ = 16.18, p < .01) while that of the far objects had been largely underestimated (t_(39)_ = -9.72, p < .01). This is visible in figure 2, by comparing the estimates for each target’s distance (near in open circles, far in filled circles) with the corresponding simulated values (1m and 7m dashed lines, respectively). As a result, the size estimates at near and far were closer to each other in magnitude than their corresponding physical values, which suggests that the constant retinal size of the target overrode much of the size estimation based on oculomotor signals. In agreement with this, the correlation between the responses in degrees and the target angles (r = .88) was significantly higher (z = 8.42, p < .01) than the correlation between the responses in cm and the target diameters (r = .67).

Overall, this pattern of results is compatible with a phenomenon previously termed “visuospatial compression”, by which the perceptual space tends to compress along the depth dimension relative to the physical space, such that near and far objects appear closer to each other than they are (Campagnoli et al., 2017). Perceived size, which is lawfully related to perceived distance, warps accordingly: For example, a small coin near the body and the moon in the sky are judged as more similar in size if viewed such that their images subtend the same area on the retina and are surrounded uninformative background, because their retinal projections tend to be scaled by a common distance (Gilinsky, 1951; Gogel, 1969). Compared to the veridical values, participants matched a larger size to the near target and a smaller size to the far target, consistent with the two stimuli appearing closer in depth and thus more similar in diameter. Figure 2 shows the signature of this effect: the near and far estimates all lie close to the dashed line representing the target’s size projected at the probe’s distance of 4 m. This perceptual bias was promoted by three experimental conditions: first, the adjustments were made at the same intermediate distance, which caused near and far images to be rescaled by a similar factor; second, the near and the far targets subtended the same exact retinal size to begin with, flattening their two estimates towards a common value; third, and most critical for the study, the overwhelming lack of depth information in the surroundings of the target under the No-Depth condition prevented the visual system from contrasting the perceptual compression of space.

The Depth condition was designed to measure the degree to which retinal depth cues allowed observers to disambiguate how far and how big the target actually was. Under this condition, in addition to ocular vergence, participants could rely on plenty of pictorial cues to assess the target’s egocentric distance. As expected, participants adjusted the near and the far targets as if they appeared more dissimilar in size relative to No-Depth (F_(1,39)_ = 233.65, p < .001), compatible with them also appearing more distant from one another.

### Explicit-egocentric versus implicit-retinotopic encoding of 3D error information

A possible explanation as to why the error encoding mechanism based on depth cues integration did not affect implicit learning is that adaptation may rely on a different, lower-level form of spatial processing, such as, for example, expressing the error size retinotopically as the angular distance between the projections of the endpoint and of the target’s center. In principle, such an encoding mechanism is effectively more stable and resistant to contextual changes, because it constructs an error signal based on local features taken out of their context. The downside of its specificity is a lack of flexibility, for example if changes in the context suddenly render those same features as irrelevant for the action’s goal.

In fact, this description of how implicit learning may function in 3D space fits well with the behavioral results of the previous studies conducted in 2D, which characterize sensorimotor adaptation as stereotyped, slow and largely independent of the context or task demands (Taylor and Ivry, 2011; Morehead et al., 2017). Notably, it also fits well with recent neuroimaging studies showing that the cerebellum contains multiple topographic maps of the visual field that are retinotopically organized (Buckner et al., 2011; van Es et al., 2019; Winawer and Curtis, 2019). These findings could suggest that the computation of an error signal is somehow expressed in retinal coordinates. Another similarity with the existing literature is that implicit learning appears to be more insensitive compared to explicit learning (Bond & Taylor, 2015, 2017; Morehead et al., 2017; Kim et al., 2018). Indeed, a retinotopic mechanism for localizing error information is computationally more parsimonious than scaling retinal projections in egocentric coordinates.

Since the encoding of a 3D scene is distributed among multiple stages in the visual system, it could be hypothesized that each learning mechanism accessed a distinct phase of the analysis. The simplest level is when explicit depth estimates have not emerged yet: In order to re-aim from a point A to a point B on a 2D surface, it is in principle sufficient to hold a retinotopic mapping of the locations of the two points without having to know their physical distance in space, akin to when making saccades. In this case, the spatial relationship between point A and point B can be expressed in terms of the visual angle between them, which is also invariant of their distance from the observer. This information, combined with few other retinal cues such as the radial expansion and compression of the optic flow, is powerful enough to support several complex tasks ranging from human navigation (Foo et al., 2005) to landing aircrafts (Gibson et al., 1955).

An egocentric-versus-retinotopic dichotomy could be the hint to a possible specialization of the motor learning components with respect to depth information, where implicit adaptation contributes maximally within the near space whereas explicit learning handles errors in the far space. Retinal vision is especially important in the peripersonal space, where things are usually manipulated. Within reachable distance, retinal disparity is by far the most reliable of the depth cues (Howard and Rogers, 1995) and most common manipulatory actions can be thought of as alignment tasks. For example, grasping a cup of coffee can be successfully done by simply nulling the relative disparity between the fingers and the cup’s surface, without retrieving their absolute location in space (Melmoth and Grant, 2006; Melmoth et al., 2007). However, since disparity decays rapidly with the square of the distance, monocular pictorial cues constitute the predominant signals for the interpretation of the 3D space beyond ∼2 meters (Howard and Rogers, 2002). The interaction between distance and visual condition found in this perceptual task is a classic signature of this competition between cues. When the stimulus was at the near distance of 1m, binocular disparity was strong enough to allow observers to detect that the target did not change in size (nor in egocentric position) between the Depth and the No-Depth scenes. Conversely, binocular disparity was negligible when the target was at the far distance of 7m, thus size (and position) estimation was entirely driven by the richness in the pictorial cues.

It is possible that sensorimotor adaptation would selectively respond to errors in the peripersonal space because small corrections that are meaningful near the body correspond to enormous overcompensations at large distances, which would require a counter-intervention by the explicit system. At far distances, the competition between the two types of learning would trigger a positive feedback loop that would not converge towards a stable state. Moreover, a near/far division of labor is consistent with previous work showing that implicit learning is most efficient with small errors that are within the range of the motor noise, whereas it saturates with larger perturbations that are instead handled by the explicit learning (Miyamoto et al., 2020).

If explicit and implicit learning were equally competitive at large distances, their interaction would be very problematic for the motor system, since even small errors would result in great instability. Furthermore, the fact that the movement kinematics of the Clamped group did not reflect changes in the apparent distance of the target whatsoever (as opposed to the Delayed group) is another evidence in favor of a true null effect of depth. Considering the many analogies between the data collected here and the previous literature, the present study hints that the implicit learning might be more concerned with the direction of a perturbation, while explicit learning with its magnitude (Bond and Taylor, 2015; Morehead et al., 2017; Butcher and Taylor, 2018; but see Kim et al., 2018). Finally, recent data showed that implicit learning could decrease its contribution over time, akin to an inhibition mechanism, when the motor system faces task demands that make sensorimotor adaptation counterproductive to the performance (Avraham et al., 2021; Albert et al., 2021).

### Conclusion

As the interaction between the two learning systems is multi-faceted and complex, this investigation has just scratched the surface of the problem. Future work will need to address how depth cues are incorporated in detail and how explicit and implicit systems work in cooperation or in competition with each other.

## Acknowledgments

We thank Cameron Hayes and Chandra Greenberg for their help with data collection.

